## Supplemental figures for "Ensuring Robustness in Scientific Research, Split-root assays as an example case"

### Supplementary material

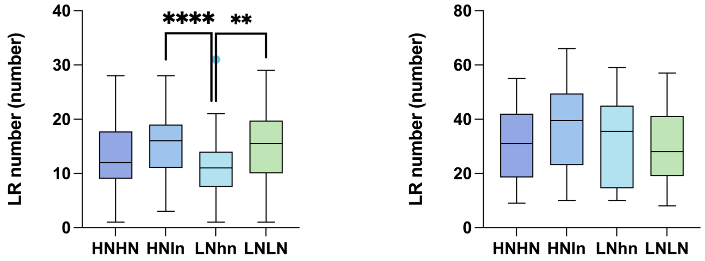


**Figure S1. Comparison of the LRs numbers of Arabidopsis split-root for different nitrogen availabilities in a short (15 DAG) and a long (20 DAG) protocol.** The Number of LR of *Arabidopsis thaliana* was measured in response to 5 days of treatment in split-root conditions in either 15 days old seedlings (left panel) or in 20 days old seedlings (right panel). LRs responses to different conditions: the high nitrogen (10mM KNO3) in both sides of the split-root condition, HNHN; the side of high nitrogen in the split-root heterogeneous condition, HNln; the side of low nitrogen in the split-root heterogeneous condition, LNhn; and the low nitrogen (0.2mM KNO3) in both sides of the split-root condition, LNLN. Boxplot displays the median of each group bounded by the first and third quartile. Asterisks indicate the significance in the comparison of two nitrogen treatments: *P<0.05; **P<0.01; ***P<0.001. The HNln vs LNhn split-root treatment was compared using ratio paired t test and the rest of the comparisons were analyzed using one-way ANOVA.

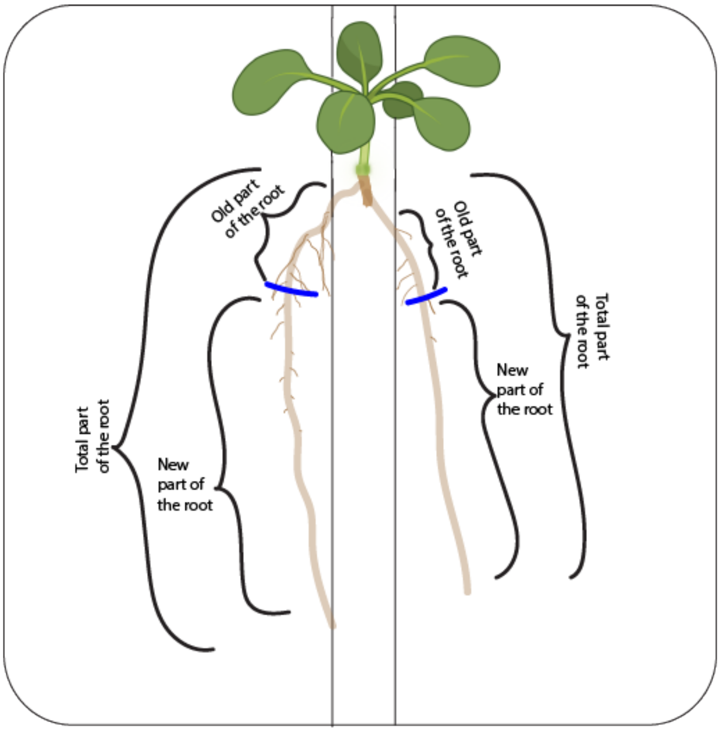


**Figure S2. Diagram of the different sections in the root analyses.** Blue lines indicate the position of the root at the start of split root treatment and are routinely used during experiments. The old part represents the section of the main root of the plant that had already developed at day 0 of the treatment when the plant was transferred to the split-root plate, and its associated LRs. The new part represents the section of the main root that developed during the treatment and its associated LRs. The total part of the roots is the sum of the old and new parts.

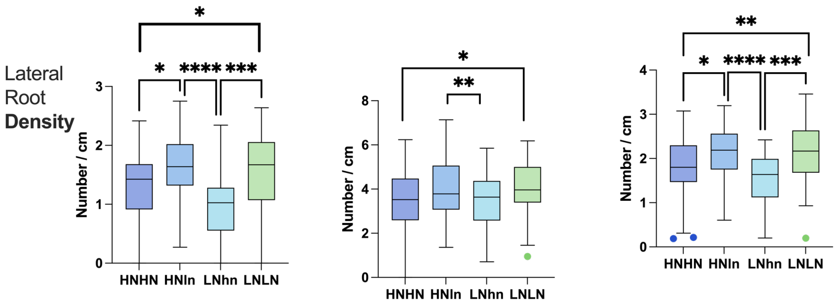


**Figure S3. Comparison of the LR density of Arabidopsis in split-root for different nitrogen availability in different sections of the root.** *Arabidopsis thaliana* root system response in 15 DAG old seedlings after 5 days of treatment in split-root conditions. LR density in response to each condition, high nitrogen (10mM KNO3) in both sides of the split-root condition (HNHN), the side of high nitrogen in the split-root heterogeneous condition (HNln), the side of low nitrogen in the split-root heterogeneous condition (LNhn), and the low nitrogen (0.2mM KNO3) in both sides of the split-root condition (LNLN). The LR density was measured for 3 sections, the new section (left panel), the old section (middle panel), and the total section (right panel). Asterisks indicate the significance in the comparison of two nitrogen treatments: *P<0.05; **P<0.01; ***P<0.001. The HNln vs LNhn split-root treatment was compared using Wilcoxon test and the rest of comparison were made using Kruskal-Wallis test.

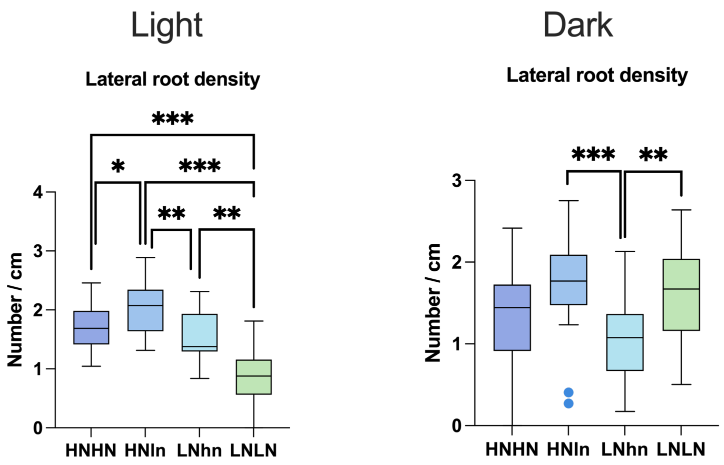


**Figure S4. Comparison of the LR density of Arabidopsis in split-root for different nitrogen availability with the roots covered or exposed to light**. *Arabidopsis thaliana* 15DAG old root system response after 5 days of treatment in split-root conditions in either seedlings with roots exposed at light (non-covered) (left) or seedlings with roots under D-root system (right). LRs responses to each condition, high nitrogen (10mM KNO3) in both sides of the split-root condition (HNHN), the side of high nitrogen in the split-root heterogeneous condition (HNln), the side of low nitrogen in the split-root heterogeneous condition (LNhn), and the low nitrogen (0.2mM KNO3) in both sides of the split-root condition (LNLN). Asterisks indicate the significance in the comparison of two nitrogen treatments: *P<0.05; **P<0.01; ***P<0.001. The HNln vs LNhn split-root treatment was compared using Wilcoxon test and the rest of comparison were made using Kruskal-Wallis test.

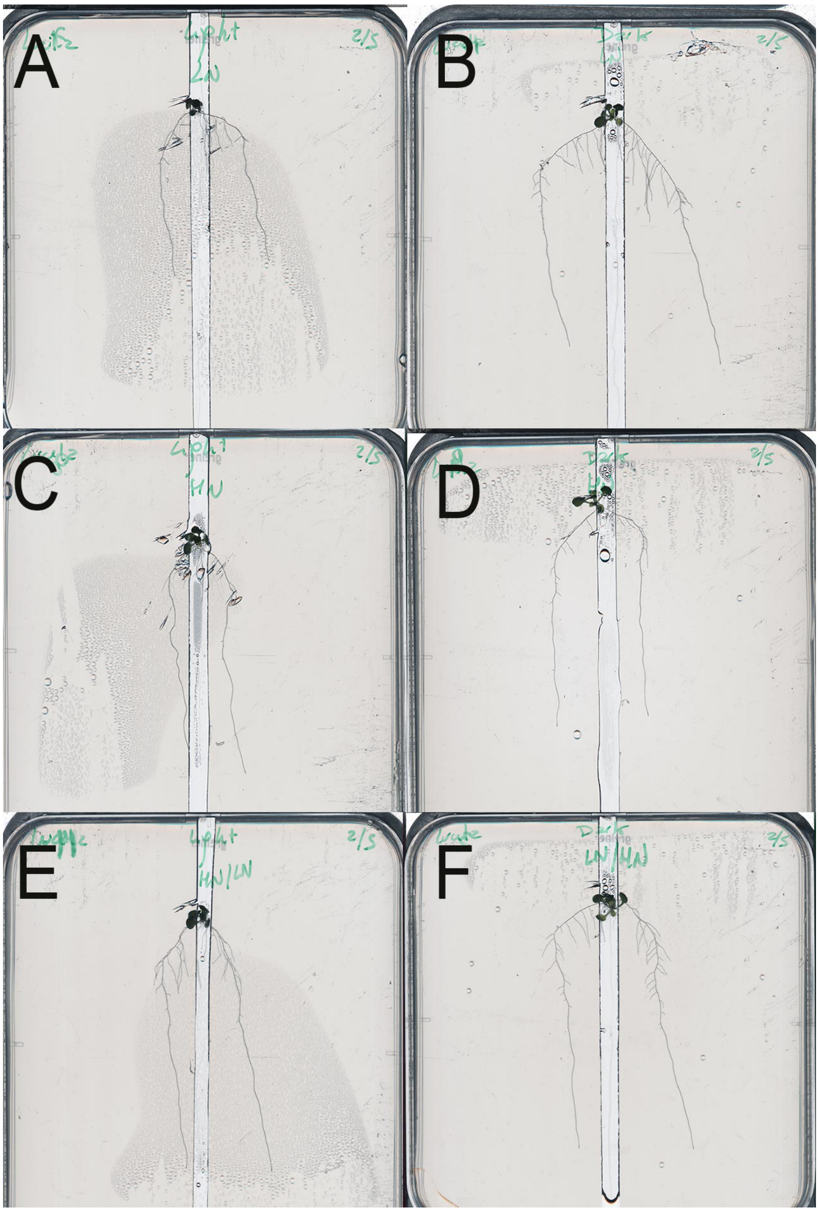


**Figure S5. Representative images of *A. thaliana* seedlings after each treatment.** *Arabidopsis thaliana* 15DAG root system response after 5 days of treatment in split-root conditions in either seedlings with roots exposed to light (non-covered) (A, C, E) or seedlings with roots grown using the D-root system (B, D, F). Seedling responses are shown to the different nitra te treatments, LNLN (A, B), HNHN (C, D), and HNln + LNhn (E, F).
