## supplemental methods for "Ensuring Robustness in Scientific Research, Split-root assays as an example case"

#### Protocol to achieve an efficient split-root assay

### Materials

| Reagent | Source | Identification Number |
|---|---|---|
| $CaCl_2$ | Sigma-Aldrich | C1016-500G |
| $MgSO_4$. $7H_2O$ | Merck | 1058860500 |
| $KH_2PO_4$ | EMSURE | 1048731000 |
| KCl | Duchefa Biochemie | P 0515 |
| $KNO_3$ | Carl Roth | 8001.2 |
| $Na_2EDTA$ (Titriplex III) | Merck | 1084181000 |
| $FeSO_4.7H_2O$ | Merck | 1039650500 |
| $MnSO_4.4H_2O$ | Merck | 1.02786 |
| $H_3BO_3$ | Fisher Scientific | 17-1322-01 |
| $ZnSO_4.7H_2O$ | Sigma-Aldrich | Z4750 |
| $Na_2MoO_4 \cdot 2H_2O$ | Sigma-Aldrich | M1651-100G |
| $CuSO_4$ | Fluka | 15633380 |
| MES hydrate | Thermo Scientific | H56472.36 |
| Plant agar | Duchefa Biochemie | P1001.1000 |
| Sucrose Crystallized | Duchefa Biochemie | S0908.5000 |
| Thick Bleach | Zone | 8714773111321 |

| Material | Brand | Specifications |
|---|---|---|
| Seeds | *Arabidopsis thaliana* Columbia-0 | Lab stocks. NASC: N1093 |
| Micropore tape | Leukopor | 1.25 cm x 9.2 m |
| Scanner | Epson | Perfection v800 Photo |
| Razor blade | Personna | Stainless Steel Double Edge Blade. 60-0139 |
| Scalpel | Swann-Morton | Carbon Steel, Number 23 |
| Tweezers | Dumont | Style: 7. sn.1091.59.2 |
| D-root system | Self-made in our workshop | |
| Plates | Greiner | 120x120x17 mm |

**Before you begin**

Split-root protocols are extensively used to study long-distance signaling *in Arabidopsis thaliana*. It provides a robust and powerful method to separate local, systemic, and long-distance signaling in response to environmental cues experienced by the roots. This variant of the method is adjusted to study young seedlings' responses to heterogenous nitrogen in between the two halves of the root and it makes use of the D-root system (Silva-Navas et al., 2015) to prevent light exposure of roots.

### Methods

#### Preparation of solid germination medium

**Timing:** 1 day

Before the experiment, prepare the medium for the sowing process. To achieve a higher germination success rate and higher homogeneity in resulting seedling sizes, a small amount of sucrose is added to the media (0.3 mM), and the initial concentration of nitrogen is chosen such that it is sufficient for the plant to grow normally, not displaying either a foraging or systemic repression response when the whole plant is grown on this concentration (10 mM). All the specifications about the nutrients and their manufacturer are described in the Materials section.

**Table 1. List of nutrients and their concentration in the germination medium.**

| Macro | Final concentration (mM) |
|---|---|
| $CaCl_2$ | 1.5 |
| $MgSO_4$ | 0.75 |
| $KH_2PO_4$ | 0.625 |
| KCl | 0 |
| $KNO_3$ | 10 |
| **Micro** | **Final concentration (µM)** |
| Fe(III)Na-EDTA | 50 |
| $MnSO_4$ | 50 |
| $H_3BO_4$ | 50 |
| $ZnSO_4$ | 15 |
| $Na_2MoO_4$ | 0.5 |
| $CuSO_4$ | 0.1 |

1. Preparation of the germination medium:
    a. Prepare the nutrient solution in the volume needed for the experiment, adding all the nutrients mentioned in **Table 1** with mQ water (start with 95% of the desired volume to allow the volume expansion from the added components).
    b. After all the nutrients have dissolved, add 1g/L of MES to the solution
    c. Supplement the media with 0.3 mM of sucrose.
    d. Adjust the pH to 5.8 with the addition of 1M KOH.
    e. Pour media to autoclavable bottles that contain 1% (w/v) agar.
    f. Autoclave the medium at 120°C for 20 minutes.
    g. Once the media has cooled down to 60°C, pour 25 mL with a pipette boy into each 12x12 cm square petri dish in sterile conditions.
    h. Let the plates air dry with the lid open.

**Note:** The plates can be used as soon as the medium has solidified. Alternatively, the plates can be stored at 4°C for 2-3 weeks.

**Optimized parameter:** The amount of sucrose added was optimized to maximize homogeneity in germination success and seedling growth while avoiding negative effects on the desired nitrogen response phenotypes. We initially tried 0 mM of sucrose, but this resulted in too large heterogeneity in germination and growth of the plants, leaving too few plants to continue the next steps of the experiment with. High sucrose concentrations (15 and 30 mM) were tried as well, yet these led to a failure to reproduce the usual nitrogen response in our settings. A sucrose concentration of 0.3mM gave us sufficient homogeneity in plant germination and growth while maintaining the desired nitrogen foraging phenotypes.

#### Preparation of the split-root plates.

**Timing:** 1 day

The preparation of the split-root plates should be done at least 24 hours in advance of the split-root transfer to allow the nutrient solutions to diffuse in the agar. Plates will be supplemented with either $KNO_3$ solution or KCl solution to achieve the desired NO3 concentration for the experiment while maintaining a constant K concentration.

**Table 2. List of nutrients and their final concentration in the treatment medium.**

| Macro | Final **concentration** (mM) |
|---|---|
| $CaCl_2$ | 1.5 |
| $MgSO_4$ | 0.75 |
| $KH_2PO_4$ | 0.625 |
| KCl | 0 |
| $KNO_3$ | 0.2 |
| Micro | Final concentration (µM) |
| Fe(III)Na-EDTA | 50 |
| $MnSO_4$ | 50 |
| $H_3BO_4$ | 50 |
| $ZnSO_4$ | 15 |
| $Na_2MoO_4$ | 0.5 |
| $CuSO_4$ | 0.1 |

1. Preparation of the treatment medium:
    a. Prepare the nutrient solution in the volume needed for the experiment, adding all the nutrients mentioned in **Table 2** to mQ water (start with 95% of the desired volume to allow the volume expansion from the added components).
    b. After all the nutrients have dissolved, add 1 g/L of MES to the solution.
    c. Supplement the media with 0.3 mM of sucrose.
    d. Adjust the pH to 5.8 with the addition of KOH.
    e. Adjust the volume to the exact desired volume with water.
    f. Pour media into autoclavable bottles that contain 1% (w/v) agar.
    g. Autoclave the medium at 120°C for 20 minutes.

h. Once the media has cooled down to 60°C, pour 25 mL of media into each 12x12 cm plate.
i. Let the plates air dry with the lid open.

2. Fabrication of the trenches.
    a. Once the plates are completely dry proceed to create a trench (i.e. make cuts and remove the agar) of approximately 0.5 cm in the middle of the plate from the top to the bottom of the plate **(Figure 1).**

**Note:** We use a custom-made guillotine **(Figure 2)** device created in our workshop facilities for this, but the cuts can also be performed with a scalpel. The chosen cutting device should be flame-sterilized before use.

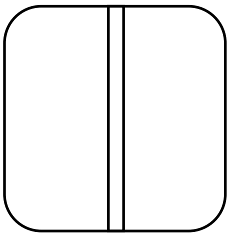

**Figure 1. Graphic illustration of where the trench is in the plate**

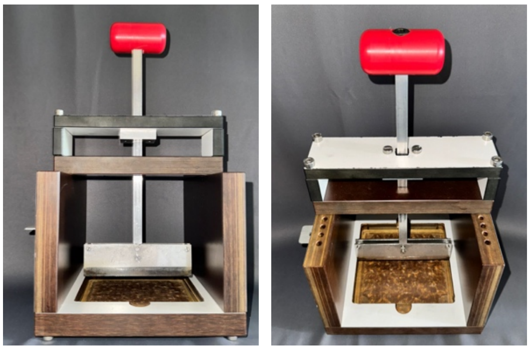

**Figure 2. Front and top photographs of the custom-made guillotine.**

b. After the cutting, remove the agar in the trench with a flame-sterilized spatula. The spatula should not be wider than the trench.

**Note:** No agar should remain in the trench, and the two blocks of agar in the plate should have become fully disconnected to avoid diffusion between different treatment compartments.

3. Preparing nutrient treatments.

**Note:** Three different types of plates will be made in this step: 1) HNHN, a homogeneous plate where both sides of the trench have a high nitrogen (HN) concentration, 2) LNLN, a homogenous plate with both sides with low nitrogen (LN) concentration, 3) and HNLN, a heterogeneous plate with a high nitrogen concentration on one side and a low nitrogen concentration on the other side of the trench.

a. Make 1mM stock solutions of $KNO_3$ and KCl each and autoclave them.

b. Pipette 118 µl of $KNO_3$ to the high nitrogen (HN) agar compartments, and 118 µl KCl to the low nitrogen agar compartments.

**Note:** Distribute the 118 µl into small droplets over the patch for a more homogenous distribution.

c. Use a sterile L-shaped cell spreader to spread the solution evenly over the patch until it is fully absorbed, and no liquid remains on the surface.
d. Close the plates and store them at 4°C.

**Note:** Plates can be stored for up to 2-3 weeks.

**Optimized parameter:** The same level of sucrose was used in both treatment and germination media, and was optimized for maximizing germination and growth homogeneity while maintaining the desired nitrogen response phenotypes.

**Optimized parameter:** The LN concentration chosen for this experiment was not random. After trying 0.5 mM and 0.2 mM of nitrate (based on previous literature) more reproducible results were found to arise from using the lowest concentration.

#### Growth of the plant material

**Timing:** 2 weeks

This section describes the procedure for growing seedlings with 2 lateral roots developing as 2 main root systems needed for the split-root treatment assay.

1. Sterilization of the seeds.

**Note:** The following describes liquid sterilization, but your preferred alternative sterilization method can also be used.

a. Place at least 10% more seeds than needed for sowing in an Eppendorf tube and add 1 mL of 70% ethanol.
b. Shake the tube until well mixed.
c. Spin the seeds down.
d. Remove as much ethanol as possible by pipetting.
e. Add 1 mL of commercial bleach (diluted 1:10) to the tube
f. Incubate the tube for 5-8 minutes while mixing it by inversion.

**Critical:** Avoid over-incubation, as it will prevent germination.

g. Spin the seeds down.
h. Remove the bleach by pipetting.
i. Add 1 mL of sterile water to the Eppendorf.
j. Shake the Eppendorf tube by inversion.
k. Spin the seeds down.

(Steps g–k: Wash)

l. Remove the water by pipetting.
m. Repeat the wash 7 times, but do not remove the water in the final wash.

**Note:** Seeds in sterile water can be stored for up to 2-3 weeks.

2. Sowing and stratification of the seeds.

   a. Using a 10 μl pipette, place the seeds on the prepared germination plates. Each plate should contain 10 seeds, evenly spaced. Due to the use of the D-root system **(Figure 3)** that will prevent light from reaching the lower half of the plates, it is important to place the seeds in the plate just above the start of the D-root covered region. Use a stencil for a consistent positioning **(Figure 4).**

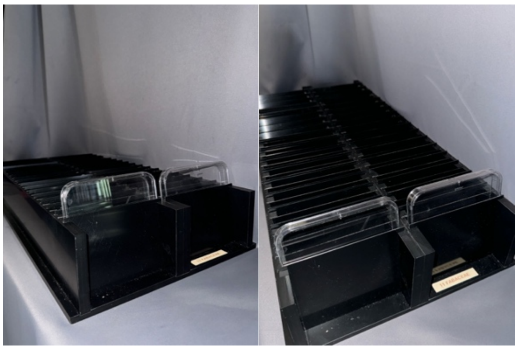

**Figure 3. Front and top photographs of the D-root system.**

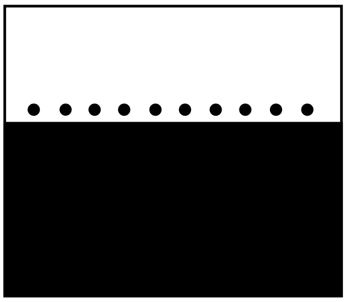

**Figure 4. Graphical illustration of the stencil used for a consistent positioning of the seeds.**

**Note: It** is recommended to sow at least 10 times more seeds than needed for the final split-root treatment assay.

   b. Tape the plates with micropore tape (to avoid contamination but allow the exchange of gases) and cover them entirely from the light e.g., using aluminum foil).
   c. Store the plates at 4°C for 3-5 days for stratification.

3. Growth of the plant material.
   a. Place the plates in a D-root system device at 21°C in a room with 150 μmol $m^{-2}$ $s^{-1}$ light intensity and a 16/8 hour photoperiod.

b. After 7 days of growth, transfer the plates to sterile conditions for the cutting procedure.
   I. Discard the seedlings with roots shorter than 1.5 cm.
   II. Under a binocular microscope, cut the seedling's root 0.4-0.5 cm below the root-shoot junction using a well sharpened scalpel.
   III. Lift the cut end of the root from the agar using tweezers.
   IV. Remove the remaining root from the plate with tweezers, minimizing agar damage.
   V. Close the plates with new micropore tape and return them to the growth chamber for 3 days of recovery.

**Note:** If there is a lateral root at the cutting point, cut below it.
**Note:** All the instruments are sterilized before use either with flame sterilization or with a heat-block at 300°C.
⚠**Critical:** Avoid damaging the agar to prevent roots from growing inside the agar instead of on its surface.
**Optimized parameter:** Our protocol is adjusted to the use of a D-root system. We demonstrated in the result section of the paper **(Figure 5)** that exposing the roots to light using this same protocol will affect the response to low nitrogen. If a D-root system is not available, another way to cover the roots from light is using aluminum foil.

#### Split-root treatment

**Timing:** 5 days

Select the most uniform seedlings from the previous step for the split-root treatment. Requirements for selection:

- Seedlings must have only two lateral roots.
- The lateral roots should not differ by more than 0.1 cm in length.
- Avoid plants with adventitious roots at the root-shoot junction.
- The length variation of the lateral roots among selected seedlings should not exceed 0.5 cm.
- Discard plants with callus formation at the cut site **(Figure 5)**.

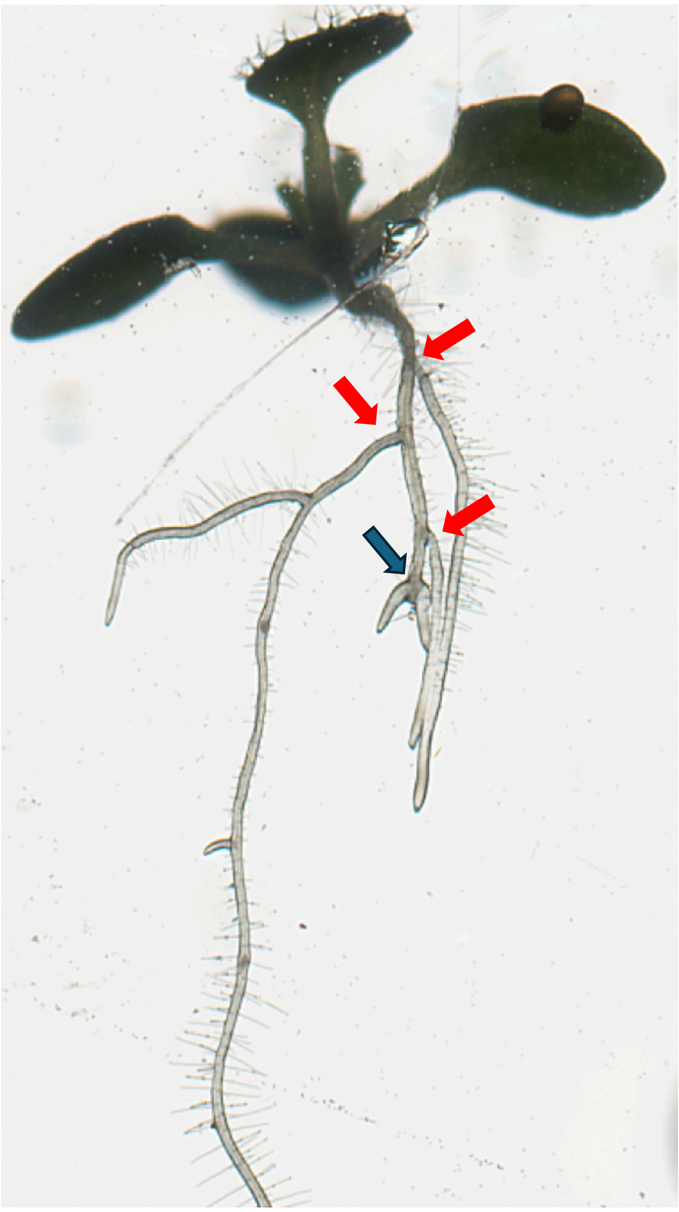

**Figure 5. Example of callus formation at the cut site.** Red arrows indicate lateral roots formed before the cut. The blue arrow indicates lateral roots that are formed at the exact point of the cut.

**Optimized parameters:** Older seedlings were used for split-root experiments (20 DAG - 10 days of growth, 5 days of recovery and 5 days of treatment) and yielded results reproducing the outcomes. We selected the younger 15 DAG seedlings for our protocol due to faster experiment times, easier handling of the plants, and easier data collection.

1. Placing the seedlings on the split-root plates.
    a. Use two tweezers to gently scoop the seedlings under both roots and the cotyledons. Place them with one root on each half of the split-root plate **(Figure 6)**. Ensure the shoot is positioned high enough to avoid shading from the D-root system. Place only one seedling per plate.

📝**Note:** Lateral roots should form an angle of 30-60° with the trench to prevent bending due to gravitropism (when angle is too large) or growing into the trench (when angle is too small).

📝**Note:** To avoid absorption by the cotyledons (MacGregor et al., 2008), place the shoot in the trench.

⚠**Critical:** Avoid touching the lateral root tips with the tweezers to prevent damage or stress to the meristems.

⚠ **Critical:** Ensure the entire root is in contact with the agar except for the primary root which should be placed in the trench.

b. Close and seal the plates with micropore tape.
c. Mark the endpoint of each lateral root with a marker on the plate, or scan them to record the starting point of the treatment.
d. Place the plates under the same growth conditions as before.
e. After 120 hours of treatment, scan the plates at 600 dpi.
f. For image processing and root system architecture (RSA) analysis, use the Smartroot plugin for ImageJ.

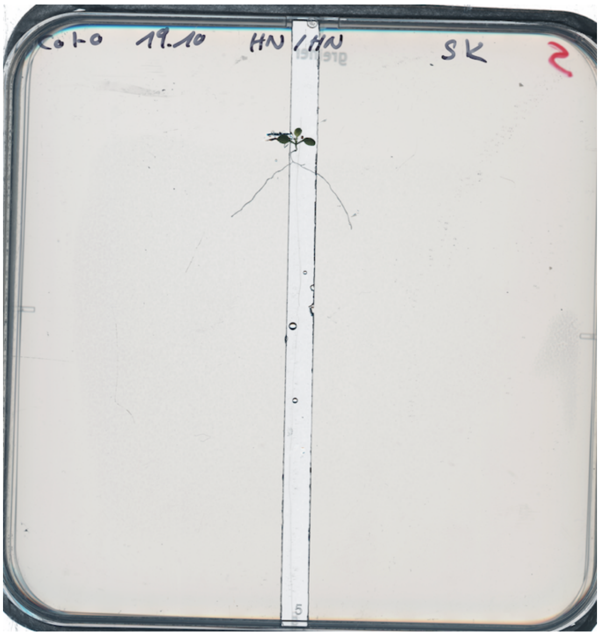


**Figure 6. Example of a seedling placed in a split-root plate.**
